# Susceptibilities of human ACE2 genetic variants in coronavirus infection

**DOI:** 10.1101/2021.07.18.452826

**Authors:** Wenlin Ren, Yunkai Zhu, Jun Lan, Hedi Chen, Yuyan Wang, Hongyang Shi, Fei Feng, Da-Yuan Chen, Brianna Close, Xiaomin Zhao, Jianping Wu, Boxue Tian, Zhenghong Yuan, Dongming Zhou, Mohsan Saeed, Xinquan Wang, Rong Zhang, Qiang Ding

**Affiliations:** Center for Infectious Disease Research, School of Medicine, Tsinghua University, Beijing 100084, China; Key Laboratory of Medical Molecular Virology (MOE/NHC/CAMS), School of Basic Medical Sciences, Shanghai Medical College, Biosafety Level 3 Laboratory, Fudan University, Shanghai 200032, China; School of Life Sciences, Tsinghua University, Beijing 100084, China; School of Pharmaceutical Sciences, Tsinghua University, Beijing 100084, China; CAS Key Laboratory of Molecular Virology and Immunology, Institut Pasteur of Shanghai, Chinese Academy of Sciences, Shanghai, China; Department of Biochemistry, Boston University School of Medicine, Boston, MA, United States of America; National Emerging Infectious Diseases Laboratories, Boston University, Boston, MA, United States of America; Key Laboratory of Structural Biology of Zhejiang Province, School of Life Sciences, Westlake University, 18 Shilongshan Road, Hangzhou 310024, Zhejiang Province, China; Department of Pathogen Biology, School of Basic Medical Sciences, Tianjin Medical University, Tianjin, China

**Keywords:** COVID-19, SARS-CoV, SARS-CoV-2, HCoV-NL63, ACE2, SNP.

## Abstract

The COVID-19 pandemic, caused by SARS-CoV-2, has resulted in more than 1603 million cases worldwide and 3.4 million deaths (as of May 2021), with varying incidences and death rates among regions/ethnicities. Human genetic variation can affect disease progression and outcome, but little is known about genetic risk factors for SARS-CoV-2 infection. The coronaviruses SARS-CoV, SARS-CoV-2 and HCoV-NL63 all utilize the human protein angiotensin-converting enzyme 2 (ACE2) as the receptor to enter cells. We hypothesized that the genetic variability in ACE2 may contribute to the variable clinical outcomes of COVID-19. To test this hypothesis, we first conducted an *in silico* investigation of single-nucleotide polymorphisms (SNPs) in the coding region of ACE2 gene. We then applied an integrated approach of genetics, biochemistry and virology to explore the capacity of select ACE2 variants to bind coronavirus spike protein and mediate viral entry. We identified the ACE2 D355N variant that restricts the spike protein-ACE2 interaction and consequently limits infection both *in vitro* and *in vivo*. In conclusion, ACE2 polymorphisms could modulate susceptibility to SARS-CoV-2, which may lead to variable disease severity.

## INTRODUCTION

Coronaviruses (CoVs) are enveloped viruses with positive-sense, single-stranded RNA genomes that belong to the *Coronaviridae* family and *Orthocornavirinae* subfamily^1, 2^. There are four genera in this subfamily: *Alphacoronavirus, Betacoronavirus, Gammacoronavirus,* and *Deltacoronavirus*^2^. In the last two decades, three betacoronaviruses have caused outbreaks of severe pneumonia in humans: the severe acute respiratory syndrome coronavirus (SARS-CoV)^3^, the Middle East respiratory syndrome coronavirus (MERS-CoV)^4^, and SARS-CoV-2^5–7^, the cause of the ongoing COVID-19 pandemic.

Similar to SARS, the severity of COVID-19 disease is positively correlated with increased age, preexisting comorbidities, and/or other environmental factors^8^. Host genetic factors could also contribute to clinical outcomes, as demonstrated for other infectious diseases such as HIV and malaria^9–12^. Given the variable incidence and disease severity of COVID-19 in different regions and ethnicities of the world, human genetic variation has attracted increasing attention for its potential role in SARS-CoV-2 transmission and pathogenicity^13, 14^. Genome-wide association studies of severe COVID-19 patients have identified several genetic risk factors, including 6 genes found in a region of chromosome 3^15, 16^. In addition, some individuals with life-threatening COVID-19 pneumonia were found to have a deficiency in their type I interferon (IFN)-mediated immune response due to genetic mutations or autoimmune antibodies targeting type I IFN^17, 18^.

SARS-CoV-2, as well as SARS-CoV and human coronavirus NL63 (HCoV-NL63), utilizes the human protein ACE2 as a cellular receptor to gain entry into human cells^19–22^. The viral spike (S) protein of NL63, SARS-CoV, and SARS-CoV-2 is comprised of an S1 and S2 domain, which are separated by a protease cleavage site. The S1 domain directly and specifically binds the peptidase domain of ACE2 via its receptor-binding domain (RBD), exposing the cleavage site for processing by host proteases. This ultimately leads to S2-mediated virus-host cell membrane fusion in a species-specific manner^23^. A number of variations have been observed in the ACE2 gene, some of which have been significantly associated with arterial hypertension, diabetes mellitus, cerebral stroke, coronary artery disease, heart septal wall thickness and ventricular hypertrophy^24^. However, it is unknown whether these natural ACE2 variants decrease or increase their affinity for coronavirus S protein and affect the susceptibility of individuals to infection.

In this study, we performed a series of biochemical and functional experiments to assess the impact of ACE2 SNPs on interaction with coronavirus S proteins and SARS-CoV-2 entry *in vitro* and *in vivo*. This led to the identification of an SNP D355N [rs961360700] that potentially protect individuals against SARS-CoV-2 infection. Our study suggest that ACE2 polymorphism may alter human susceptibility to SARS-CoV-2 infection, and contribute to ethnic and geographical differences in SARS-CoV-2 spread.

## RESULTS

### Genetic polymorphisms in the human ACE2 gene could impact interaction with coronavirus spike proteins

ACE2 is a peptidase that is expressed on the surface of many cell types, including lung epithelial cells, and regulates the renin-angiotensin-aldosterone system^25^. ACE2 polymorphisms associated with hypertension (HT) and diabetic heart disease have been previously described^26, 27^. Here we surveyed the Genome Aggregation Consortium Database (gnomAD) (https://gnomad.broadinstitute.org/) and found that human ACE2 is highly polymorphic, with 223 ACE2 single-nucleotide variants (SNVs) that result in missense mutations. We hypothesized that these human ACE2 SNVs could influence susceptibility to SARS-CoV, SARS-CoV-2, and/or HCoV-NL63 and potentially affect disease outcomes. To test this, we chose 12 SNVs that led to amino acid substitutions at or in close proximity to the interface of the SARS-CoV, SARS-CoV-2 or HCoV-NL63 S protein complexed with human ACE2 (**Fig. 1A-B**) ^28–32^. Of the 13 ACE2 SNVs, 4 have a mutated residue at the interface with the SARS-CoV-2 or SARS-CoV S protein (T17A [rs781255386], E37K [rs146676783], M82I [rs766996587] and D355N [rs961360700])^28–30, 32^ and 2 at the interface with the NL-63 S protein (E37K [rs146676783] and D355N [rs961360700])^31^.

**Figure 1.**
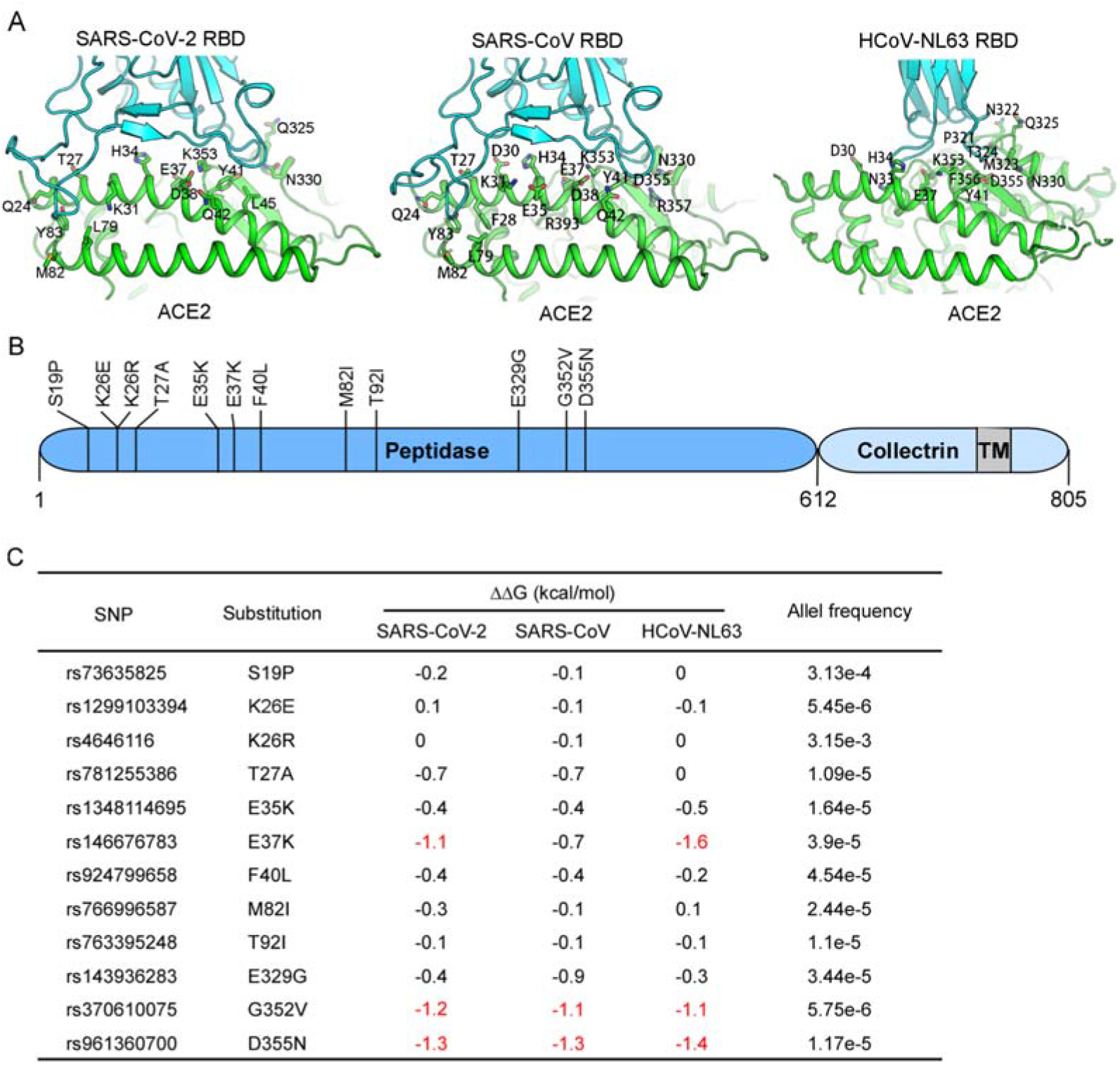
Schematic representation of the ACE2 molecule and positions of the studied SNP loci. (**A**) The structures of human ACE2 complexed with the spike proteins of SARS-CoV-2 (PDB code: 6M0J), SARS-CoV (PDB code: 2AJF) or HCoV-NL63 (PDB code: 3KBH). ACE2 and the spike protein of each virus are colored in green and cyan, respectively. The residues of ACE2 at the interface with each spike protein are highlighted. (**B**) Coding-region variants from gnomAD in the genes encoding ACE2 used in this study are indicated. The SNPs and the alteration of the amino acids in this study are shown. (**C**) Prediction of the interaction of coronavirus spike proteins with ACE2 variants. The ΔΔG for missense mutation was calculated by mCSM-PPI2 with PBD 2AJF (SARS-CoV spike complexed with human ACE2), 6M0J (SARS-CoV-2 spike complexed with human ACE2) or 3KBH (NL63-CoV spike complexed with human ACE2) as the model. The individual SNPs are named according to their identification numbers registered at the SNP database (dbSNP). The allele frequency of SNPs are as referenced in gnomAD database.

To elucidate the potential impact of these mutations on viral entry, we predicted the binding affinity of the ACE2 variants and the viral spike proteins. By using the computational platform mCSM-PPI2^33^, we assessed the difference between the binding Gibbs free energy (ΔG) of the wild-type ACE2/spike protein complex with that of each variant ACE2/spike protein complex, thus calculating the ΔΔG_wild type–mutant_ value. According to the mCSM-PPI2 calibration, ΔΔG >+1 kcal/mol is a confident indication that a variant has enhanced binding to the viral spike protein relative to the wild-type protein whereas a ΔΔG <-1 kcal/mol indicates a variant has impaired binding. Based on these criteria, most of the ACE2 SNVs exhibited limited alteration in their interaction with the viral spike proteins compared to wild-type ACE2 (**Fig. 1C**). However, the SNV E37K was predicted to have a significantly decreased interaction with the SARS-CoV-2 spike (ΔΔG=-1.1 kcal/mol) and even more so with the NL63 spike (ΔΔG=-1.6 kcal/mol). G352V and D355N showed dramatically reduced binding with all three spike proteins. The other variants were considered to have an uncertain impact on binding (**Fig. 1C**).

### Human ACE2 variants bind coronavirus spike proteins with varying efficiencies in a cell-based assay

We next employed a cell-based assay that used flow cytometry to assess the binding of the viral S proteins to human ACE2 variants^34, 35^. We cloned the cDNA of each human ACE2 variant (mouse ACE2 was included as a negative control) with a C-terminal FLAG tag into a bicistronic lentiviral vector, pLVX-IRES-zsGreen1. Since this vector contained the fluorescent protein zsGreen1 cloned under an internal ribosomal entry site (IRES) element, we used zsGreen1 expression as a measure of transduction efficiency. We transduced HeLa cells, which lack endogenous ACE2 expression^19^, and performed fluorescence-activated cell sorting (FACS) to collect cells with comparable expression levels of the ACE2 variants (**Fig. S1A and B**).

Next, we incubated HeLa-ACE2 variant cells with the purified fusion protein consisting of the S1 domain of the coronavirus S proteins being examined (SARS-CoV-2, SARS-CoV or HCoV-NL63) and an Fc domain of human lgG (S1-Fc) (1 μg/ml). Binding of the fusion proteins to ACE2 was quantified by flow cytometry and the binding efficiency was calculated as the percent of S1-Fc positive cells among ACE2-expressing cells (zsGreen1+) (**Fig. 2A**). As expected, the S1-Fc proteins of SARS-CoV-2 or NL63 did not bind to HeLa cells expressing mouse ACE2 and showed levels comparable to that of the empty vector control, whereas SARS-CoV S1-Fc exhibited 21% binding efficiency.

**Figure 2.**
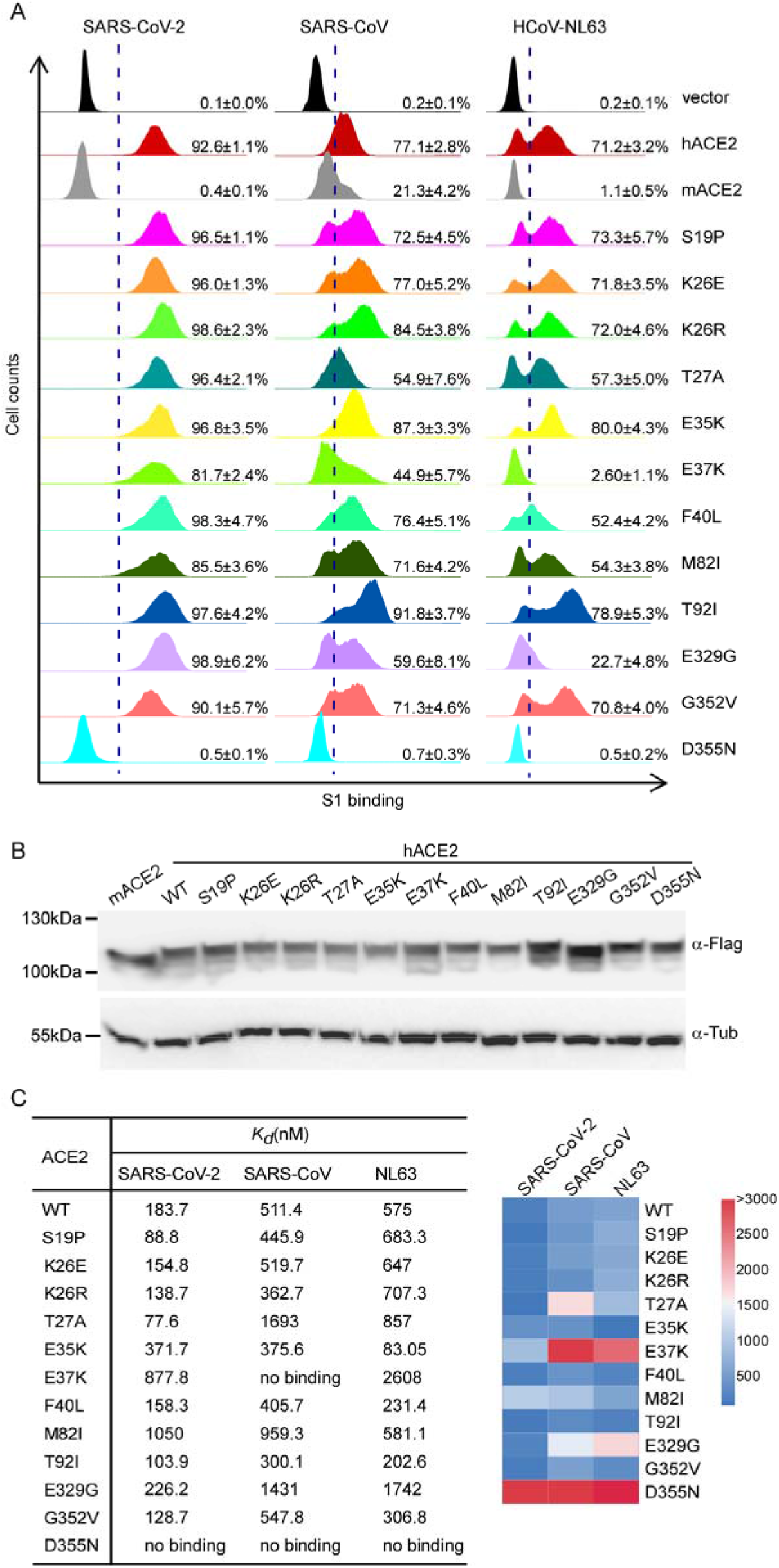
ACE2 variants bind viral spike proteins. (**A**) HeLa cells transduced with ACE2 variants were incubated with the recombinant S1 domain of the SARS-CoV-2, SARS-CoV or HCoV-NL63 spike proteins C-terminally fused with Fc (1μg/ml). Cells were then incubated with goat anti-human IgG (H+L) conjugated to Alexa Fluor 647 followed by flow cytometry analysis. Values are binding efficiencies defined as the percent of ACE2-expressing cells (zsGreen1+) positive for S1-Fc. Error bars represent the SD of the mean from one representative experiment with three biological replicate samples and this experiment was independently repeated three times. (**B**) HeLa cells transduced with lentiviruses expressing FLAG-tagged human ACE2 variants were subjected to immunoblotting. Tubulin served as the loading control. This experiment was independently repeated three times with similar results. A representative blot is shown. (**C**) Affinity of ACE2 SNVs for coronavirus spike proteins. Dissociation constant (K_d_) values were determined by SPR and are presented as a heatmap according to the indicated color legend. This experiment was independently repeated three times with similar result and the results from one representative experiment are shown.

For HeLa cells expressing the human ACE2 variants, the S1-Fc proteins exhibited varying levels of binding efficiencies. For example, SARS-CoV-2 S1-Fc bound to ACE2 variants at a level similar to that of wild-type ACE2 (range of 82%-98% vs 93%), with the exception of D355N variant that showed dramatically impaired binding (0.5%). Interestingly, SARS-CoV S1-Fc protein bound to some ACE2 variants (K26R, E35K, T92I) at a significantly higher efficiency (84.5%, 87.3%, 91.8%, respectively) compared to the wild-type ACE2 (77.1%). In contrast, other variants, such as T27A, E37K, and D355N showed decreased binding affinity, with the latter being the most impaired variant (0.7%).

Of the three coronavirus S proteins tested, the one from HCoV-NL63 bound human ACE2 the least efficiently (71.2%). In addition, it exhibited similar binding efficiencies for most of the ACE2 variants, except for E37K, E329G and D355N. While E329G bound 3-fold less efficiently than wild-type ACE2 (22.7% vs 71.2%), E37K and D355N were almost completely impaired in binding HCoV-NL63 spike protein (2.6% and 0.5%, respectively).

Collectively, our results showed that ACE2 variants exhibited varying binding abilities with the SARS-CoV-2, SARS-CoV and HCoV-NL63 spike proteins. Some ACE2 variants had virus-specific differences in binding spike protein. D355N was severely impaired in binding all three spike proteins tested. Increasing the concentration of SARS-CoV-2 S1-Fc, but not that of SARS-CoV or HCoV-NL63, did allow for some detection of binding with D355N (5μg/ml, 7%; 10μg/ml, 25.7%) (**Fig. S3A, B and C**). We confirmed that the striking differences in the binding between the ACE2 variants and spike proteins were not due to differing protein expression levels or cell-surface localization (**Fig. 2B, Fig S1, Fig. S2 and Fig. S4**). Together, these observations suggest that despite using the same cellular receptor, these coronaviruses evolved to utilize distinct amino acids of ACE2 for cell entry.

### Binding of ACE2 SNVs with coronavirus spike protein *in vitro* by SPR analysis

To further assess the binding of the ACE2 SNVs with the coronavirus spike proteins, we expressed and purified recombinant ACE2 variants, SARS-CoV-2 receptor-binding domain (RBD), SARS-CoV RBD, and HCoV-NL63 RBD, and directly assayed the protein binding *in vitro* by surface plasmon resonance (SPR) analysis (**Fig. S5A-B**). The dissociation constant (K_d_) for wild-type ACE2 binding the SARS-CoV-2 RBD was 183.7nM while that of E37K or M82I was about 5-6 fold higher (877.8nM and 1050nM, respectively). In line with our cell-based assay, no binding of D355N with SARS-CoV-2 was detected. The K_d_ of other SNVs with the SARS-CoV-2 RBD ranged from 77-371.7nM (**Fig. S6A, Fig. 2C**) and were consistent with the binding efficiencies determined in our cell-based assay (**Fig. 2A**).

For SARS-CoV RBD, the K_d_ for wild-type ACE2 was 511.4nM while the binding of E37K or D355N was not detected. The K_d_ for other SNVs ranged from 300-1693nM, not strikingly different from wild-type ACE2 (**Fig. S6B, Fig. 2C**). With respect to binding the HCoV-NL63 RBD, wild-type human ACE2 had a K_d_ of 575nM while E37K and E329G had 5- and 3-fold less affinity (K_d_ of 2608nM and 1742nM, respectively). Similar to the other RBD proteins, HCoV-NL63-CoV RBD did not bind the D355N variant (**Fig. S6C, Fig. 2C**).

Collectively, our SPR analysis suggests that the ACE2 SNVs possess distinct binding affinities for the three coronavirus RBDs examined. Notably, D355N bound all three coronavirus RBDs with limited affinity in both cell-based and SPR assays.

### Ability of ACE2 SNVs to mediate entry of virus pseudotyped with coronavirus spike protein

To assess the functionality of the human ACE2 variants, we evaluated their ability to support the entry of virus pseudotyped with either SARS-CoV-2 or SARS-CoV spike protein. To this end, we produced pseudotyped virus particles containing a firefly luciferase reporter gene and expressing on their surface either vesicular stomatitis virus glycoprotein (VSV-G; positive control) or the spike proteins of SARS-CoV-2 or SARS-CoV. HeLa cells expressing the ACE2 variants were then inoculated with these pseudoparticles and at 48h post-inoculation, the cells were lysed and the luciferase activity was monitored as a measure of virus entry. As expected, the VSV-G pseudoparticles readily infected cells independent of which ACE2 variant was expressed (**Fig. 3A-B, red columns**). Compared to vector-transduced HeLa cells, expression of human ACE2 enhanced the entry of SARS-CoV-2 pseudoparticles by 173-fold (**Fig. 3A-B, blue columns**). Most of the ACE2 SNVs mediated SARS-CoV-2 pseudoparticle entry at comparable levels, with the luciferase activity 125-269 fold higher than the negative control. However, viral entry was dramatically compromised in cells expressing E329G and D355N, with the luciferase activity increased by only 30- and 10-fold of the negative control, respectively.

**Figure 3.**
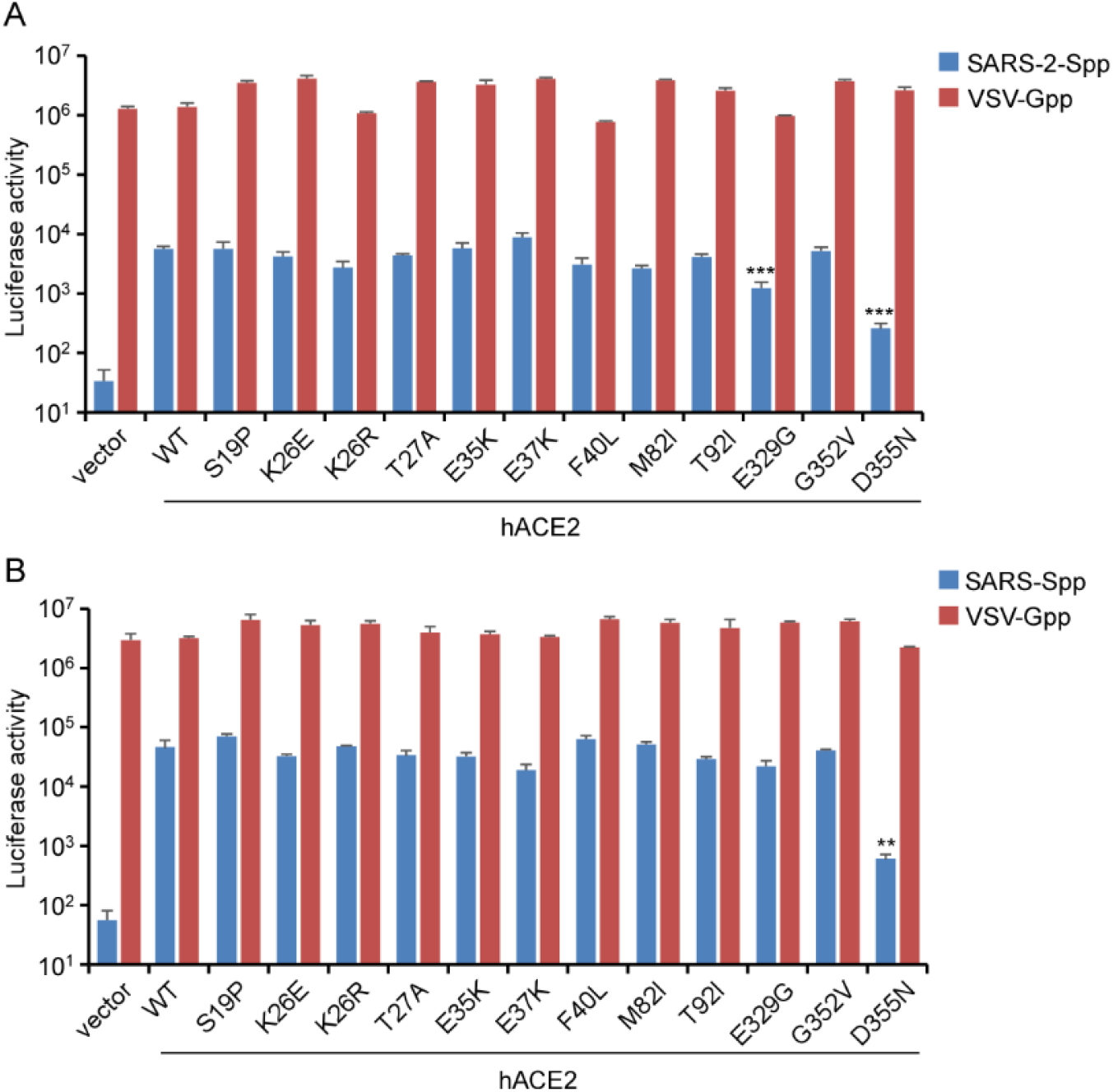
The ability of ACE2 variants to facilitate entry of virus pseudotyped with coronavirus spike proteins. (**A-B**) HeLa cells transduced with lentiviruses expressing ACE2 orthologs, SNVs or empty vector were infected with virus pseudotyped with SARS-CoV spike, SARS-CoV-2 spike or VSV-G and containing a firefly luciferase (Fluc) reporter gene. Intracellular Fluc activity of cell lysates was determined after 72h of infection. Error bars represent the SD of the mean from one representative experiment with three biological replicates and this experiment was independently repeated at least three times. **, p < 0.01, ***, p < 0.005. Significance assessed by one-way ANOVA.

Human ACE2 could mediate SARS-CoV pseudoparticle entry, as demonstrated by a 1000-fold increase in luciferase activity. Most of the ACE2 SNVs mediated SARS-CoV pseudoparticle entry to a similar extent (700-1200-fold increase over negative control). However, the luciferase activity of cells expressing D355N was severely impaired and only 32-fold higher than the negative control.

Taken together, these findings show that the ACE2 SNVs vary in their ability to support viral entry. In agreement with the data from our other assays, the D355N variant was especially limited in its ability to support SARS and SARS-CoV-2 pseudoparticle entry.

### Ability of ACE2 variants to mediate authentic SARS-CoV-2 infection *in vitro* and *in vivo*

As demonstrated above, most of the ACE2 variants were able to bind SARS-CoV-2 spike protein and support efficient entry of pseudovirus particles. However, three of the variants showed suboptimal activity against either all or a subset of the three viruses we tested. Also, some discrepancies were noted between the protein binding and pseudovirus entry assays. For example, E37K was impaired in binding with the SARS-CoV spike protein but mediated efficient entry of SARS-CoV pseudoparticles. In contrast, although E329G showed high binding efficiency for the SARS-CoV-2 spike protein, it supported SARS-CoV-2 pseudoparticle entry significantly less than wild-type ACE2. D355N was highly deficient in interacting with spike of all viruses, both in the binding as well as pseudovirus entry assays (**Fig. 2A and C**).

To further assess E37K, E329G, and D355N variants, we tested their ability to mediate authentic SARS-CoV-2 entry. HeLa cells ectopically expressing individual ACE2 variants were infected with SARS-CoV-2 virus at different doses (MOI=0.03, 0.1, 0.3 and 1) (**Fig. 4A**). At 48 h post-infection, the variant-expressing cells were fixed and stained with an antibody directed against the viral nucleocapsid (N) protein, an indicator of virus infection and replication. As expected, HeLa cells expressing mouse ACE2 were not susceptible to SARS-CoV-2 infection while those expressing wild-type human ACE2 showed high levels of infection. ACE2 SNVs E37K and E329G were comparable to wild-type ACE2 in mediating viral infection, whereas D355N supported significantly lower infection (**Fig. 4B**).

**Figure 4.**
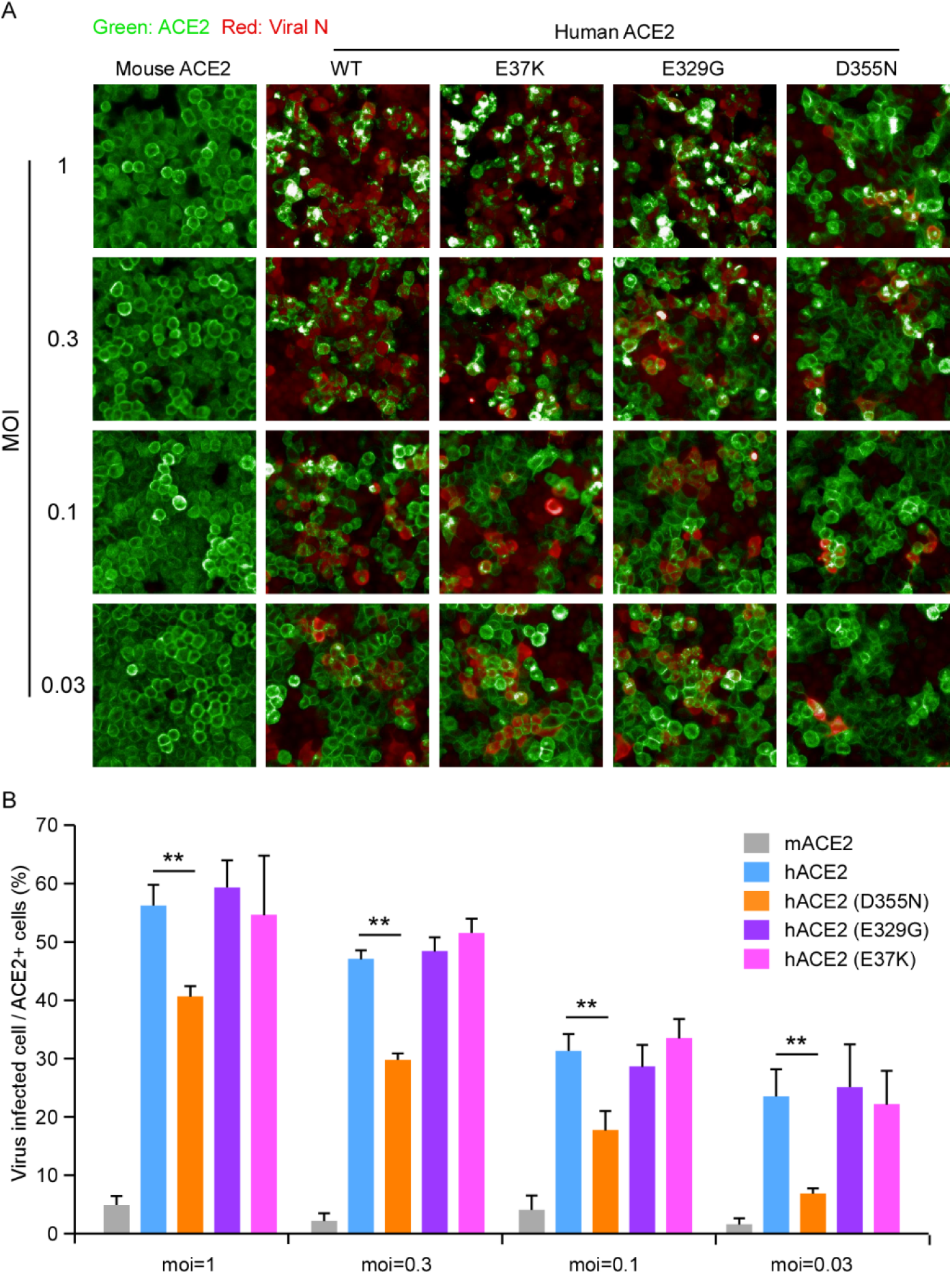
The ACE2 variants mediate authentic SARS-CoV-2 virus infection *in vitro*. **(A)** HeLa cells transduced with lentiviruses expressing human ACE2 SNVs or mouse ACE2 were infected with varying doses of SARS-CoV-2 virus (MOI=1, 0.3, 0.1 or 0.03). Expression of the viral nucleocapsid (N) protein or ACE2 orthologs was visualized by using the Operetta High Content Imaging System (PerkinElmer). Viral N protein (red) and ACE2 variant/ortholog (green) are shown. This experiment was independently repeated three times with similar results and representative images are shown. (**B**) The images were analyzed and quantified using PerkinElmer Harmony high-content analysis software 4.9. The infection efficiency represents the percentage of SARS-CoV-2 infected cells/ACE2 positive cells (y axis). Error bars represent the SD of the mean from one representative experiment with three biological replicate samples and this experiment was independently repeated three times. ns, no significance; **, p < 0.01. Significance assessed by one-way ANOVA.

Next, we further evaluated D355N *in vivo* using a mouse model transduced with replication-defective adenovirus encoding a functional human ACE2 gene which has previously been shown to support productive SARS-CoV-2 infection^34, 36, 37^. BALB/c mice were transduced intranasally with recombinant adenovirus expressing human ACE2, the D355N SNV or a vector control followed by intranasal infection with SARS-CoV-2. After 3 days of SARS-CoV-2 infection, mice were sacrificed and lung tissues collected for viral antigen detection and viral load titration by focus-forming assay (**Fig. 5A**). To examine the viral antigen spread in the lungs, we performed immunochemical staining with anti-N antibody. Viral N antigen was only well detected in lungs from infected mice transduced with hACE2 and its D355N variant, however, the viral N antigen was less abundant in that of D355N variants (**Fig. 5B**). Consistent with the N antigen staining, the greatest levels of infectious SARS-CoV-2 virus (about 1×10^6^ FFU/g of lung tissue) were in lung tissue homogenates from mice transduced with hACE2; viral load in that of D355N transduced mice was moderately reduced (about 1×10^5^ FFU/g of lung tissue), whereas virtually none or minimal levels were detected in that of mice transduced with vector (**Fig. 5C**). Taken together, our results demonstrate that D355N is limited in susceptibility to SARS-CoV-2 infection *in vitro* and *in vivo*.

**Figure 5.**
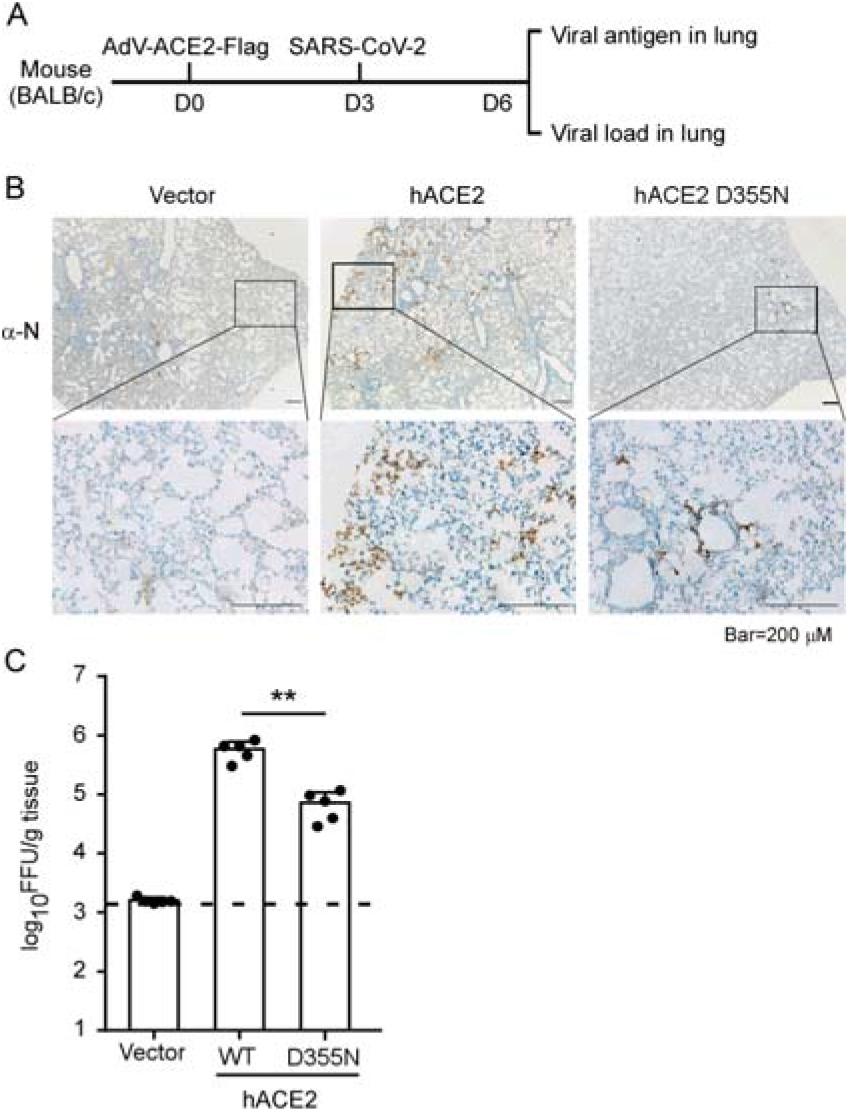
The ability of ACE2 variants to mediate authentic virus entry *in vivo*. (**A**) Schematic representation of the experimental timeline. Wild-type BALB/c mice were transduced with recombinant adenovirus expressing wild-type human ACE2 or the D355N variant ACE2 variants for 3 days, followed by SARS-CoV-2 challenge. Mice were sacrificed at day 3 post infection (n=5 mice per group) and lung tissues were collected for immunostaining with anti-N serum (**B**), and viral load titration (**C**). Representative images are shown from n = 5 mice. Scale bar, 200_μ_m (**B**). Viral load was determined by focus-forming assay. **, p < 0.01. Significance assessed by one-way ANOVA (**C**).

## DISCUSSION

Clinical outcomes of SARS-CoV-2 infection range from asymptomatic or mildly symptomatic infections to severe pneumonia, respiratory failure, and even death^23^. Epidemiological studies have identified at least three risk factors for severe disease: being male, of advanced age, and having certain co-morbidities^5, 38^. Genetic factors have also been linked to the severity of COVID-19, such as polymorphisms in a cluster of genes on chromosome 3 (including the *SLC6A20, LZTFL1, CCR9, FYCO1, CXCR6*, and *XCR1* genes)^15, 16^ and genetic deficiency in the type I IFN pathway^17^. Such findings are of utmost biological and medical importance for understanding the pathogenesis of COVID-19 and development of countermeasures. ACE2 is the receptor for SARS-CoV, SARS-CoV-2 and HCoV-NL63^19, 21, 22, 39^ and is also the major genetic determinant of host range and tissue tropism for these viruses^34, 40–42^. *ACE2* encodes for a metallopeptidase that catalyzes the conversion of angiotensin II to angiotensin, which acts as a vasodilator and exerts important modulatory effects on the cardiovascular system^27^. ACE2 SNPs have been found in association with multiple disorders, including essential hypertension, dyslipidemia, hypertrophic cardiomyopathy, and ventricular hypertrophy^24^. Therefore, an investigation of the impact of ACE2 polymorphisms on SARS-CoV-2 infection is critical and could pave the way for personalized treatment strategies for COVID-19.

In this study, we evaluated how select ACE2 SNVs compare to the wild-type protein in their binding efficiency with the spike proteins of SARS-CoV, SARS-CoV-2 and NL63-HCoV. We also examined the functionality of these ACE2 variants in mediating infection of virus particles pseudotyped with these different coronavirus spike proteins as well as their ability to support authentic virus infection both *in vitro* and *in vivo*. ACE2 is one of the most polymorphous genes in human populations, with 223 missense SNVs recorded in the gnomAD database (https://gnomad.broadinstitute.org/). As the structure of viral spike protein complexed with human ACE2 has been solved^28–30^, the residues at the interface of the complex are well characterized. We chose 12 SNVs with substitutions at or closest to the complex interface, hypothesizing that these SNVs would have the greatest impact on the spike-ACE2 interaction. We identified several missense ACE2 SNVs that showed significantly altered interaction with the spike proteins of three coronaviruses we tested. The variant D355N [rs961360700] was of particular interest, as it had a significant increase in predicted binding free energy (**Fig.1B**), limited binding affinity with spikes in both cell-based and SPR assays (**Fig. 2A and C**), and reduced susceptibility to authentic SARS-CoV-2 infection *in vitro* and *in vivo* (**Fig. 4 and Fig. 5**). These findings provide experimental evidence that a human ACE2 D355N [rs961360700] polymorphism can affect susceptibility to SARS-CoV-2, SARS-CoV or HCoV-NL63 infection. Based on analysis of the structure of ACE2 complexed with SARS-CoV-2 RBD^28–30^, the loop in a beta sheet (K353-R357) of ACE2 is critical for RBD binding, and the D355N mutation could disrupt the beta sheet structure, which might explain why D355N mutant is refractory to binding by the SARS-CoV-2 spike protein.

We note that our conclusions are based on experiments performed in cell culture and in an artificial animal model and therefore have certain limitations. It will be important to extend this work to a real-world study, which could bridge the gap between genotype, phenotype and epidemiology. In addition, we only considered the effect of ACE2 SNVs. As the serine protease TMPRSS2 can prime the viral spike for direct fusion with the plasma membrane^39, 43^, SNPs in TMPRSS2 might also impact SARS-CoV-2 cell entry and be useful target of future studies. Moreover, SNPs in the 5’UTR or other non-coding regions of ACE2 were not included in this study and may also be important in regulating ACE2 transcription or translation, leading to varied ACE2 protein levels *in vivo* that could affect virus entry and the severity of COVID-19. Due to these limitations, we urge caution not to over interpret the results of this study.

Of note, SARS-CoV-2 has evolved rapidly in humans, and a variety of genomic changes, including mutations and deletions in the spike protein, have recently been identified^44, 45^. Given that genetic variants will continue to arise, it is likely that we will see more genomic changes in the spike protein that might affect the spike interaction with human ACE2. Thus, it will be important to continue testing the interactions of these mutated spike proteins with the ACE2 SNVs. In summary, our study suggest that ACE2 polymorphism could impact human susceptibility to SARS-CoV-2 infection, which contribute to ethnic and geographical differences in SARS-CoV-2 spread and pathogenicity.

## Acknowledgements

We would like to acknowledge Di Qu, Zhiping Sun, Wendong Han, Gaowei Hu and other colleagues at the Biosafety Level 3 Laboratory of Fudan University for help with experiment design and technical assistance. We thank Dr. Jenna M. Gaska for suggestions and revision of the manuscript.

This work was supported by the National Natural Science Foundation of China (32070153 to QD; 32041005 to RZ), Beijing Municipal Natural Science Foundation (M21001 to QD), Tsinghua-Peking University Center of Life Sciences (045-61020100120), Start-up Foundation of Tsinghua University (53332101319), Shanghai Municipal Science and Technology Major Project (20431900400) and Project of Novel Coronavirus Research of Fudan University.

## Materials and Methods

### Cell cultures and SARS-CoV-2 virus

HEK293T cells (American Tissue Culture Collection, ATCC, Manassas, VA, CRL-3216), Vero E6 (Cell Bank of the Chinese Academy of Sciences, Shanghai, China) and HeLa (ATCC #CCL-2) were maintained in Dulbecco’s modified Eagle medium (DMEM) (Gibco, NY, USA) supplemented with 10% (vol/vol) fetal bovine serum (FBS), 10mM HEPES, 1mM sodium pyruvate, 1×non-essential amino acids, and 50 IU/ml penicillin/streptomycin in a humidified 5% (vol/vol) CO2 incubator at 37°C. Cells were tested routinely and found to be free of mycoplasma contamination. The SARS-CoV-2 strain nCoV-SH01 (GenBank accession no. MT121215) was isolated from a COVID-19 patient and propagated in Vero E6 cells for use. All experiments involving virus infections were performed in the biosafety level 3 facility of Fudan University following the appropriate regulations.

### Plasmids

The cDNAs encoding ACE2 orthologs (Table S1) were synthesized by GenScript and cloned into pLVX-IRES-zsGreen1 vectors (Catalog No. 632187, Clontech Laboratories, Inc) with a C-terminal FLAG tag. ACE2 mutants were generated by Quikchange (Stratagene) site-directed mutagenesis. All of the constructs were verified by Sanger sequencing.

### Lentivirus production

VSV-G–pseudotyped lentiviruses expressing ACE2 orthologs tagged with FLAG at the C-terminus were produced by transient co-transfection of the third-generation packaging plasmids pMD2G (Addgene #12259) and psPAX2 (Addgene

#12260) and the transfer vector with VigoFect DNA transfection reagent (Vigorous) into HEK293T cells. The medium was changed 12 h post transfection. Supernatants were collected at 24 and 48h after transfection, pooled, passed through a 0.45-µm filter, and frozen at -80°C.

### Protein expression and purification

The RBDs of SARS-CoV-2, SARS-CoV, NL63-CoV and the N-terminal peptidase domain of wild-type and human ACE2 SNVs were expressed using the Bac-to-Bac baculovirus system (Invitrogen). Specifically, SARS-CoV-2 RBD (residues Thr333-Pro527), SARS-CoV RBD (residues Arg306–Leu515), or NL63-CoV RBD (residues Gln481-Ile616) with a N-terminal gp67 signal peptide for secretion and a C-terminal 6×His tag for purification was inserted into pFastBac-Dual vector (Invitrogen). The recombinant baculoviruses were generated according to the manufacture’s instruction to infect Hi5 cells at a density of 2×10^6^ cells/ml. After 60h, the supernatant of cell culture containing the RBD was collected, concentrated and buffer-exchanged to HBS (10 mM HEPES, pH 7.2, 150 mM NaCl). The recombinant RBD was captured by Ni-NTA resin (GE Healthcare) and eluted with 500 mM imidazole in HBS buffer. RBD was then purified by gel-filtration chromatography using the Superdex 200 column (GE Healthcare) pre-equilibrated with HBS buffer. Fractions containing RBD were collected for further analysis. The N-terminal peptidase domain of each human ACE2 (residues Met1-Asp615) variant was expressed and purified by essentially the same protocol used for RBD.

### Surface ACE2 binding assay

HeLa cells were transduced with lentiviruses expressing the ACE2 from different species for 48 h. The cells were collected with TrypLE (Thermo #12605010) and washed twice with cold PBS. Live cells were incubated with the recombinant proteins, S1 domain of SARS-CoV-2, SARS-CoV or NL63 spike C-terminally fused with Fc (1 μg/ml) at 4 °C for 30 min. After washing, cells were stained with goat anti-human IgG (H + L) conjugated to Alexa Fluor 647 (Thermo #A21445, 2 μg/ml) for 30 min at 4 °C. Cells were then washed twice and subjected to flow cytometry analysis (Thermo, Attune™ NxT).

### Surface Plasmon Resonance (SPR) experiments

SARS-CoV RBD, SARS-CoV-2 RBD or NL63 RBD was immobilized to a CM5 sensorchip (GE Healthcare) using a Biacore T200 (GE Healthcare) and a running buffer composed of 10 mM HEPES pH 7.2, 150 mM NaCl and 0.05% Tween 20. Serial dilutions of ACE2 SNVs proteins were flowed through with a concentration ranging from 1600 to 25 nM. The resulting data were fit to a 1:1 binding model using Biacore Evaluation Software (GE Healthcare).

### Production of SARS-CoV-2 or SARS-CoV pseudotyped virus

Pseudoviruses were produced in HEK293T cells by co-transfecting the retroviral vector pTG-MLV-Fluc, pTG-MLV-Gag-pol, and pcDNA3.1 expressing the SARS-CoV spike, SARS-CoV-2 spike or VSV-G (pMD2.G (Addgene #12259)) using VigoFect (Vigorous Biotechnology). At 48 h post transfection, the cell culture medium was collected for centrifugation at 3500 rpm for 10 min, and then the supernatant was subsequently aliquoted and stored at -80°C for further use. Virus entry was assessed by transduction of pseudoviruses in cells expressing ACE2 variants in 48-well plates. After 48 h, intracellular luciferase activity was determined using the Luciferase Assay System (Promega, Cat. #E1500) according to the manufacturer’s instructions. Luminescence was recorded on a GloMax Discover System (Promega).

### Immunofluorescence staining of viral nucleocapsids

HeLa cells were transduced with lentiviruses expressing the ACE2 from different species for 48 h. Cells were then infected with nCoV-SH01 at an MOI of 1 for 1 h, washed three times with PBS, and incubated in 2% FBS culture medium for 48 h for viral antigen staining. Cells were fixed with 4% paraformaldehyde in PBS, permeablized with 0.2% Triton X-100, and incubated with the rabbit polyclonal antibody against SARS-CoV nucleocapsid protein (Rockland, 200-401-A50, 1 μg/ml) at 4 °C overnight. After three washes, cells were incubated with the secondary goat anti-rabbit antibody conjugated to Alexa Fluor 488 (Thermo #A11034, 2 μg/ml) for 2 h at room temperature, followed by staining with 4’,6-diamidino-2-phenylindole (DAPI). Images were collected using an EVOS™ Microscope M5000 Imaging System (Thermo #AMF5000). Images were processed using the ImageJ program (http://rsb.info.nih.gov/ij/).

### Generation and production of recombinant adenovirus expressing ACE2 variants

cDNA of ACE2 variants with a FLAG tag at the C-terminus was cloned into pShuttle. The pShuttle-ACE2 plasmids were then electroporated 440 into BJ5183 AD-1 cells (Agilent), which were pretransformed with pAdEasy-1 to facilitate recombination with the pShuttle-CMV vector. The adenovirus constructs were then transfected into HEK293 cells. The transfected HEK293 cells were maintained until the cells exhibited complete cytopathic effect (CPE) and then harvested and freeze-thawed. The supernatants were serially passaged two more times, with harvest at complete CPE and freeze-thaw. For virus purification, the cell pellets were purified using cesium chloride density-gradient ultracentrifugation, and the number of virus particles was determined using a Nanodrop 2000 (Thermo Fisher Scientific). The adenovirus stocks were aliquoted and stored at -80°C.

### SARS-CoV-2 infection of adenovirus transduced mice

Six to eight week-old male mice (BALB/c) were transduced intranasally with adenovirus expressing wild-type ACE2, the D355N variant or empty control (5×10^10^ viral particles per mouse). After 3 days, mice were infected intranasally with SARS-CoV-2 (8×10^4^ FFU per mouse) and sacrificed at day 3 post infection. The lung tissues were harvested for histopathological analysis and virus titration. This animal experiment protocol was approved by the Animal Ethics Committee of the School of Basic Medical Sciences at Fudan University.

### Histological analysis

Lung tissues were harvested and fixed in 4% paraformaldehyde (PFA) for 48 h. Tissues were embedded in paraffin for sectioning. To detect the expression of FLAG-tagged ACE2 delivered by adenovirus, the sections were incubated in blocking reagent and then with FLAG M2 antibody (1:100 dilution, Sigma-Aldrich #1804) at 4 °C overnight, followed by incubation with HRP-conjugated goat anti-mouse IgG secondary antibody (1:5000 dilution, Invitrogen). For viral antigen detection, the sections were incubated with house-made mouse anti-SARS-CoV-2 nucleocapsid protein serum (1:5000) and HRP465 conjugated goat anti-mouse IgG secondary antibody (1:5000 dilution, Invitrogen). The lung sections from the vector-transduced mouse were used as negative control. The sections were observed under microscope (Olympus, Tokyo, Japan).

### Virus load determination by focus-forming assay

Vero E6 monolayer in 96-well plates were inoculated with serially diluted virus for 2 h and then overlaid with methylcellulose for 48 h. Cells were fixed with 4% paraformaldehyde in PBS for 1 h and permeablized with 0.2% Triton X-100 for 1 h. Cells were stained with homemade mouse anti-SARS-CoV-2 N serum overnight at 4°C, incubated with the secondary goat anti-mouse HRP-conjugated antibody for 2 h at room temperature. The focus-forming unit was developed using TrueBlue substrate (Sera Care #5510–0030).

### Statistics analysis

One-way analysis of variance (ANOVA) with Tukey’s honestly significant difference (HSD) test was used to test for statistical significance of the differences between the different group parameters. *P* values of less than 0.05 were considered statistically significant.

## Supplemental Figures and legends

**Supplemental Figure 1.**
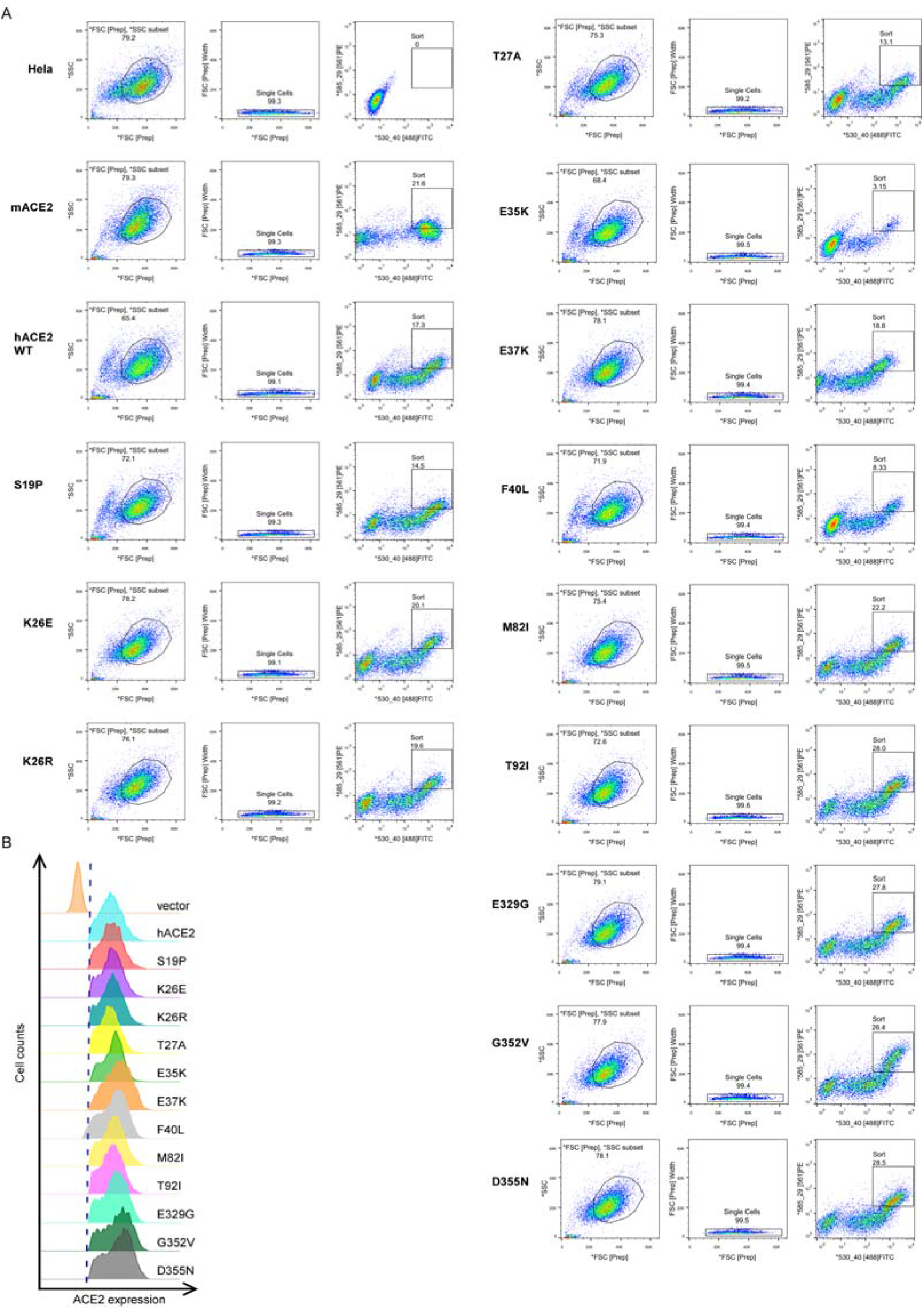
Sorting of the HeLa cells expressing comparable levels of ACE2 variants on the cell surface by fluorescence-activated cell sorting (FACS). (**A**) HeLa cells transduced with lentivirus (pLVX-IRES-zsGreen1) expressing ACE2 variants were incubated with rabbit polyclonal antibody (Sino Biological Inc. China, Cat: 10108-T24) against ACE2. The cells were washed and then stained with 2 μg/mL goat anti-rabbit IgG (H+L) conjugated to Alexa Fluor 568. The samples were subjected to FACS to sort the cells expressing comparable levels of the ACE2 variants on their cell surface. (**B**) The sorted cells were cultured and the cell surface ACE2 expression was validated again by cell surface staining as mentioned above (**A**). Cell surface ACE2 was calculated as the percent of Alex Fluor 568 positive cells among the zsGreen1-positive cells. This experiment was repeated twice with similar results.

**Supplemental Figure 2.**
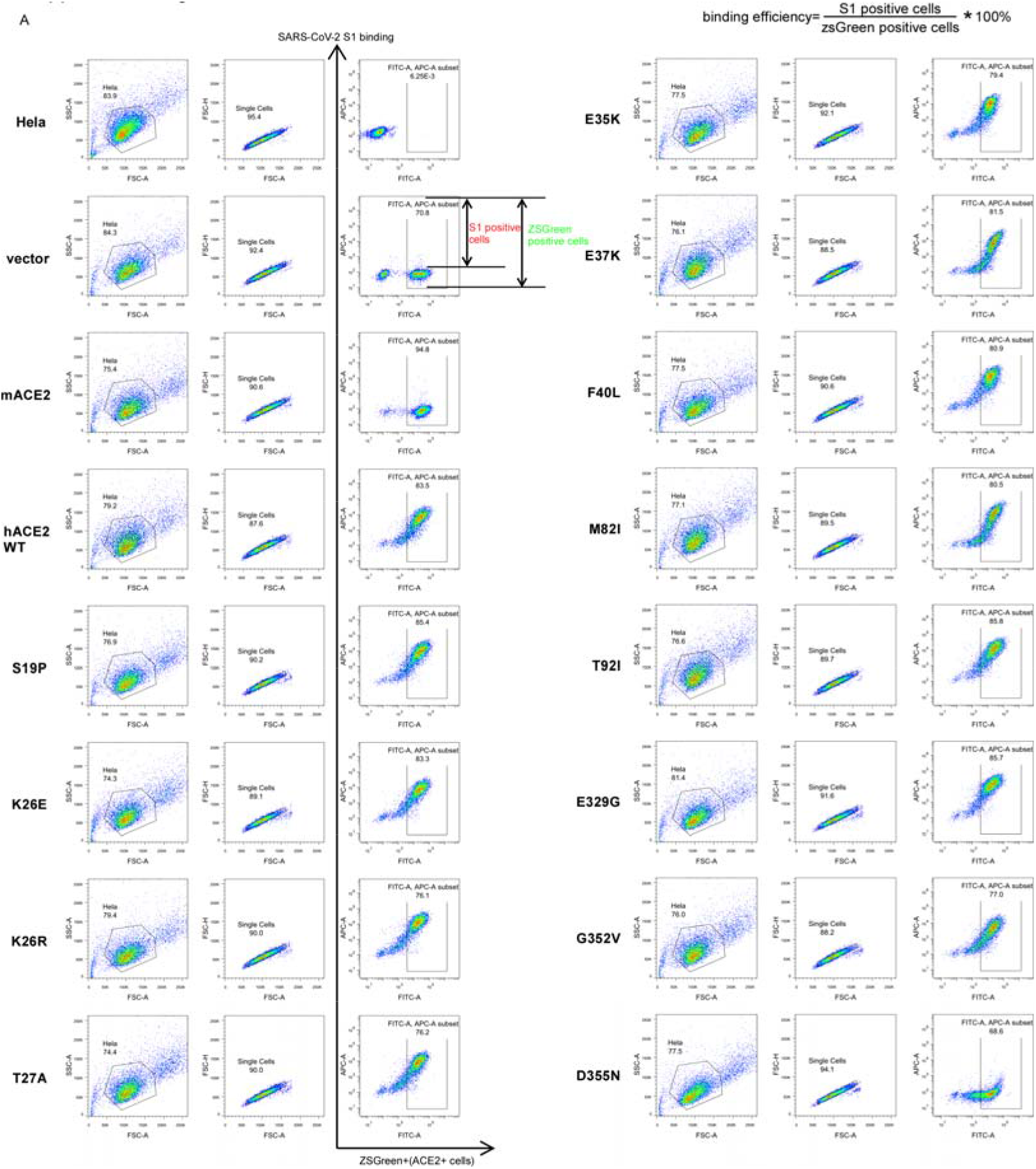
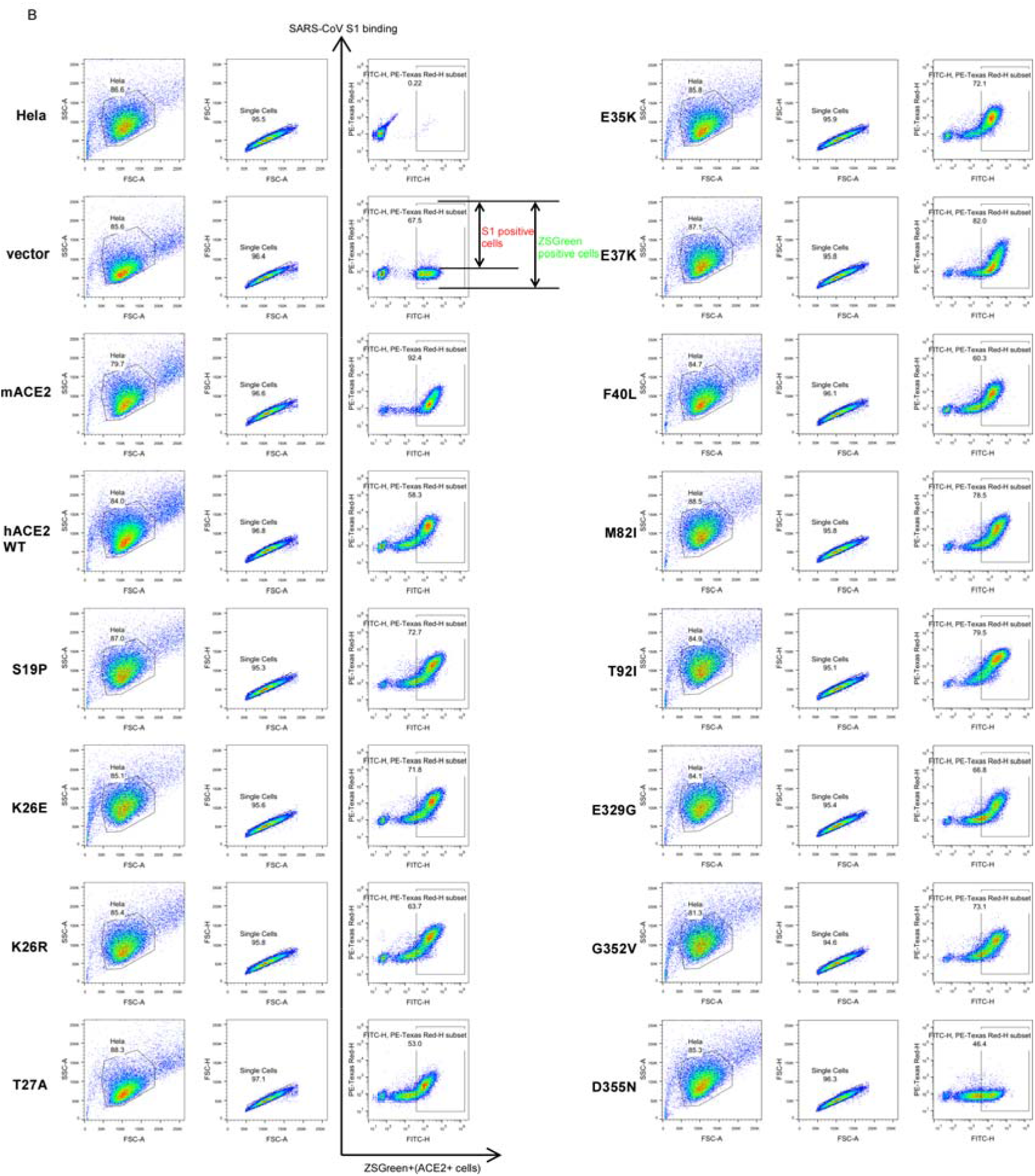
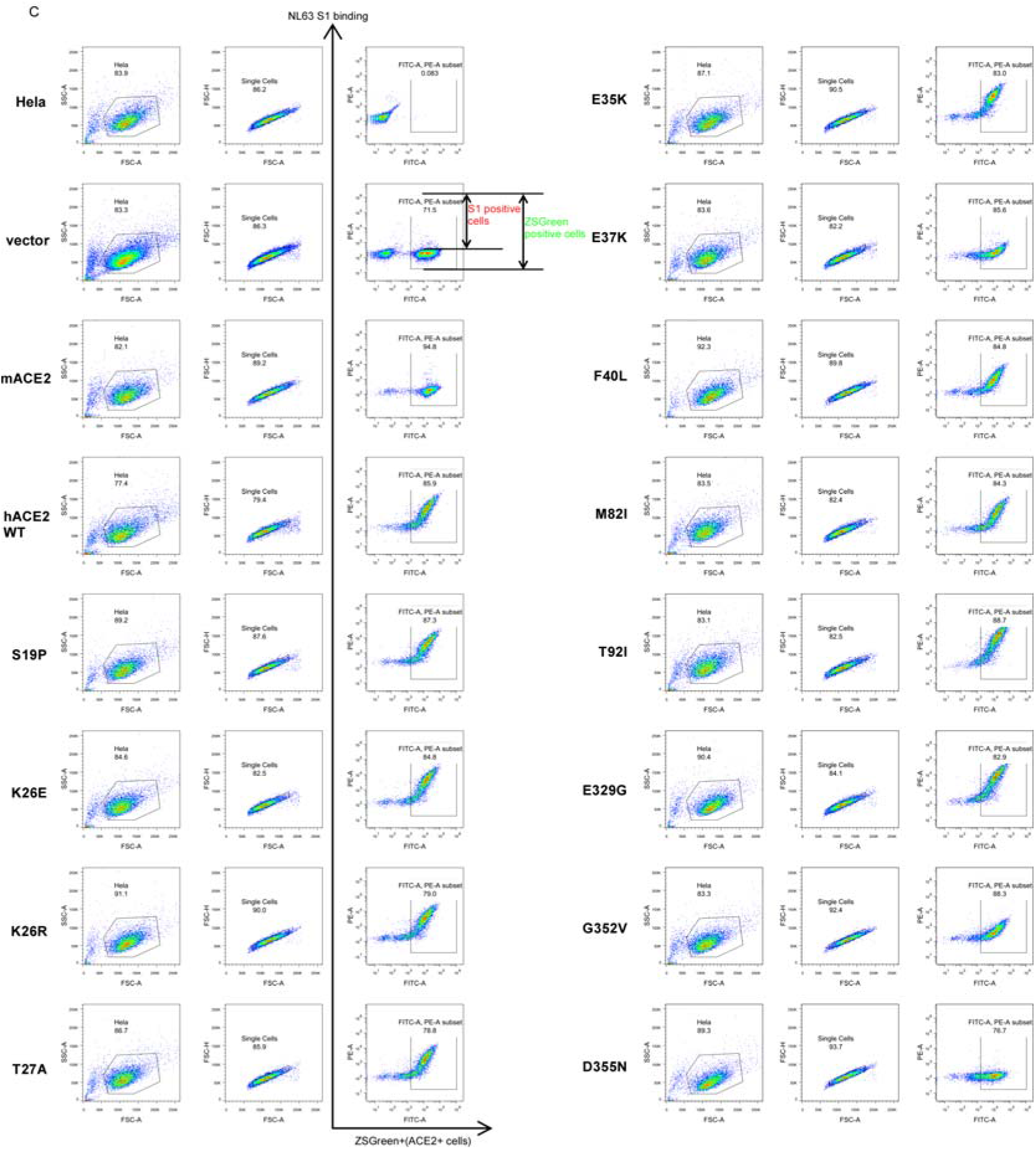
**Gating strategy for determining the binding efficiency of the ACE2 variants with SARS-CoV-2, SARS-CoV or HCoV-NL63 S1-Fc protein**. The main cell population was identified and gated on forward- and side-scatter. Single cells were further gated on FSC-A and FSC-H. The gated cells were plotted by FITC-A (zsGreen1, as the ACE2-expressing population) and APC-A (S1-Fc bound population). The FITC-A positive cell population was plotted as a histogram to show the SARS-CoV-2 (**A**), SARS-CoV (**B**) or HCoV-NL63 (**C**) S1-Fc positive population as **Fig. 2A**. The binding efficiency was defined as the percent of S1-Fc binding cells among the zsGreen1-positive cells. Shown are FACS plots representative of those used for the calculation of binding efficiencies of ACE2 variants with S1-Fc. A representative experiment is shown and this experiment was independently repeated three times with similar results.

**Supplemental Figure 3.**
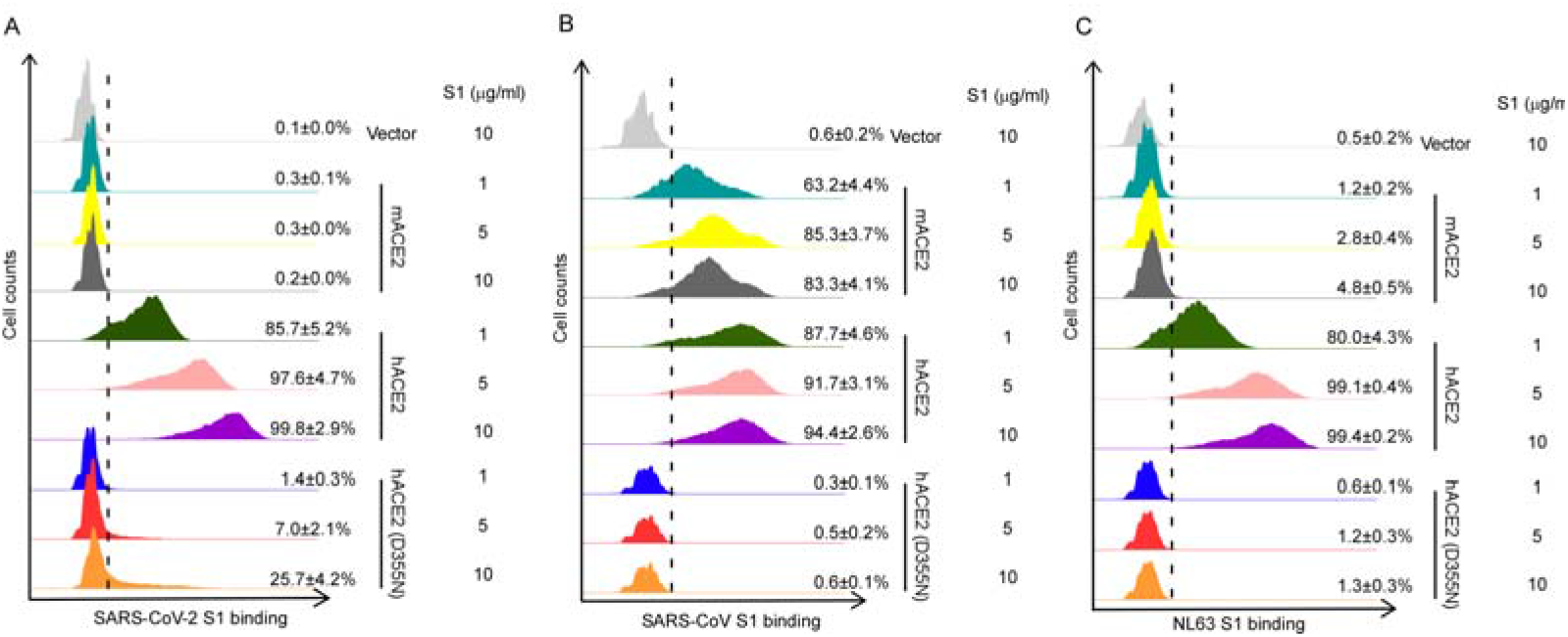
ACE2 variants bind viral spike proteins. HeLa cells transduced with ACE2 variants, were incubated with increasing doses (1μg/ml, 5μg/ml or 10μg/ml) of the recombinant S1 domain of SARS-CoV-2 (**A**), SARS-CoV (**B**) or HCoV-NL63 (**C**) spike proteins C-terminally fused to Fc with and then were stained with goat anti-human IgG (H + L) conjugated to Alexa Fluor 647 for flow cytometry analysis. Values are binding efficiencies defined as the percent of cells positive for S1-Fc among the ACE2-expressing cells (zsGreen1+ cells). Error bars represent the SD of the mean from one representative experiment with three biological replicate samples and this experiment was independently repeated three times.

**Supplemental Figure 4.**
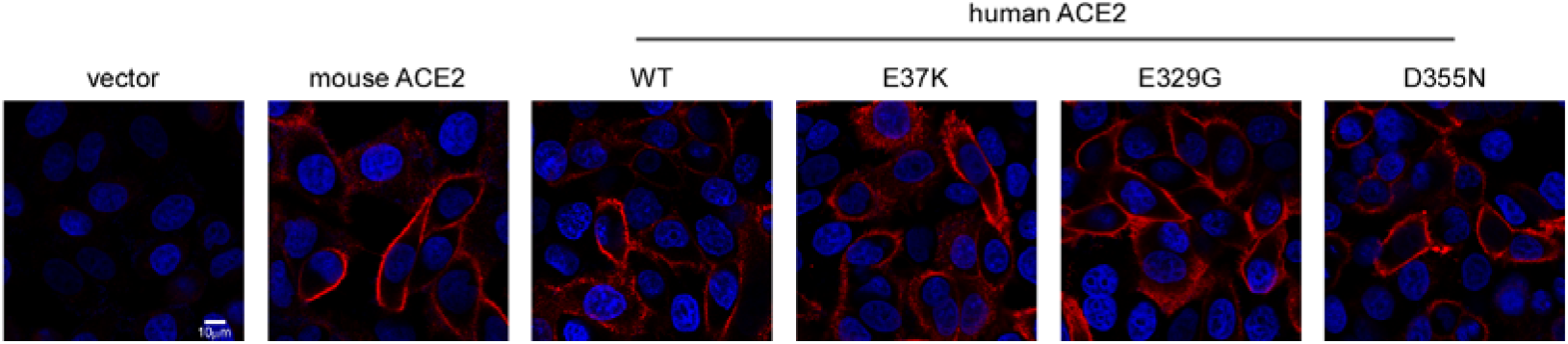
**Cell surface localization of ACE2 variants**. HeLa cells transduced with lentiviruses (pLVX-IRES-zsGreen1) expressing ACE2 variants as indicated were incubated with rabbit polyclonal antibody (Sino Biological Inc. China, Cat: 10108-T24) against ACE2. The cells were washed and then stained with 2μg/mL goat anti-rabbit IgG (H+L) conjugated with Alexa Fluor 568 and DAPI (1μg/ml). The cell images were captured with a Zeiss LSM 880 Confocal Microscope. ACE2 on cell surface was shown in the merge images processed by ZEN3.2 software. This experiment was independently repeated three times with similar results and the representative images are shown.

**Supplemental Figure 5.**
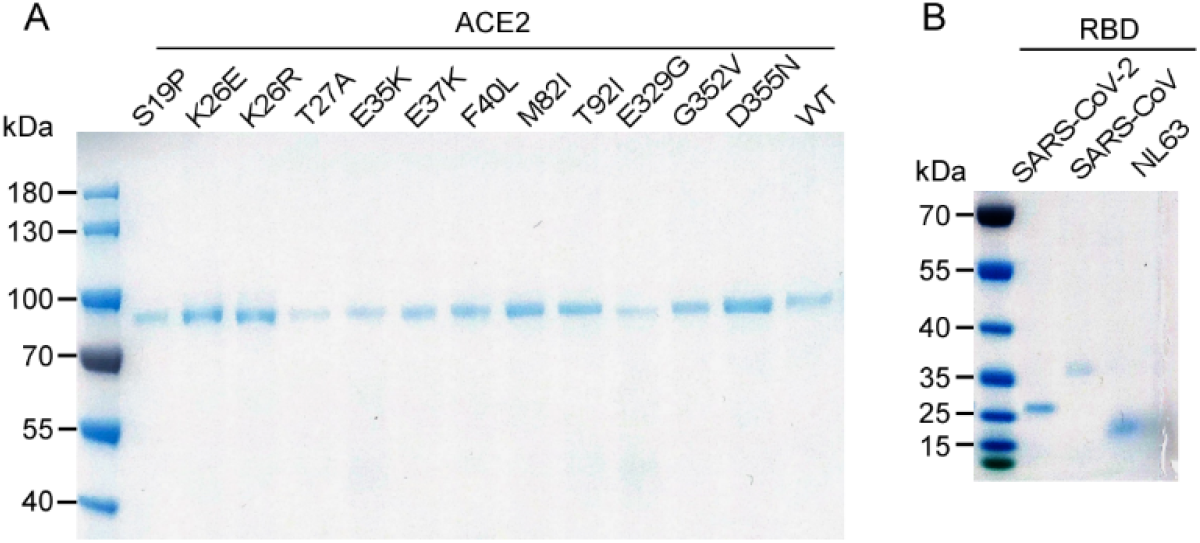
Analysis of the purified ACE2 variants and coronavirus spike proteins by SDS-PAGE. The N-terminal peptidase domain of each human ACE2 (residues Met1-Asp615) variant (**A**), SARS-CoV-2 RBD (residues Thr333-Pro527), SARS-CoV RBD (residues Arg306–Leu515), or NL63-CoV RBD (residues Gln481-Ile616) were expressed and purified as described in the ***Materials and Methods***. The purified proteins were analyzed by SDS-PAGE with Coomassie blue staining.

**Supplemental Figure 6.**
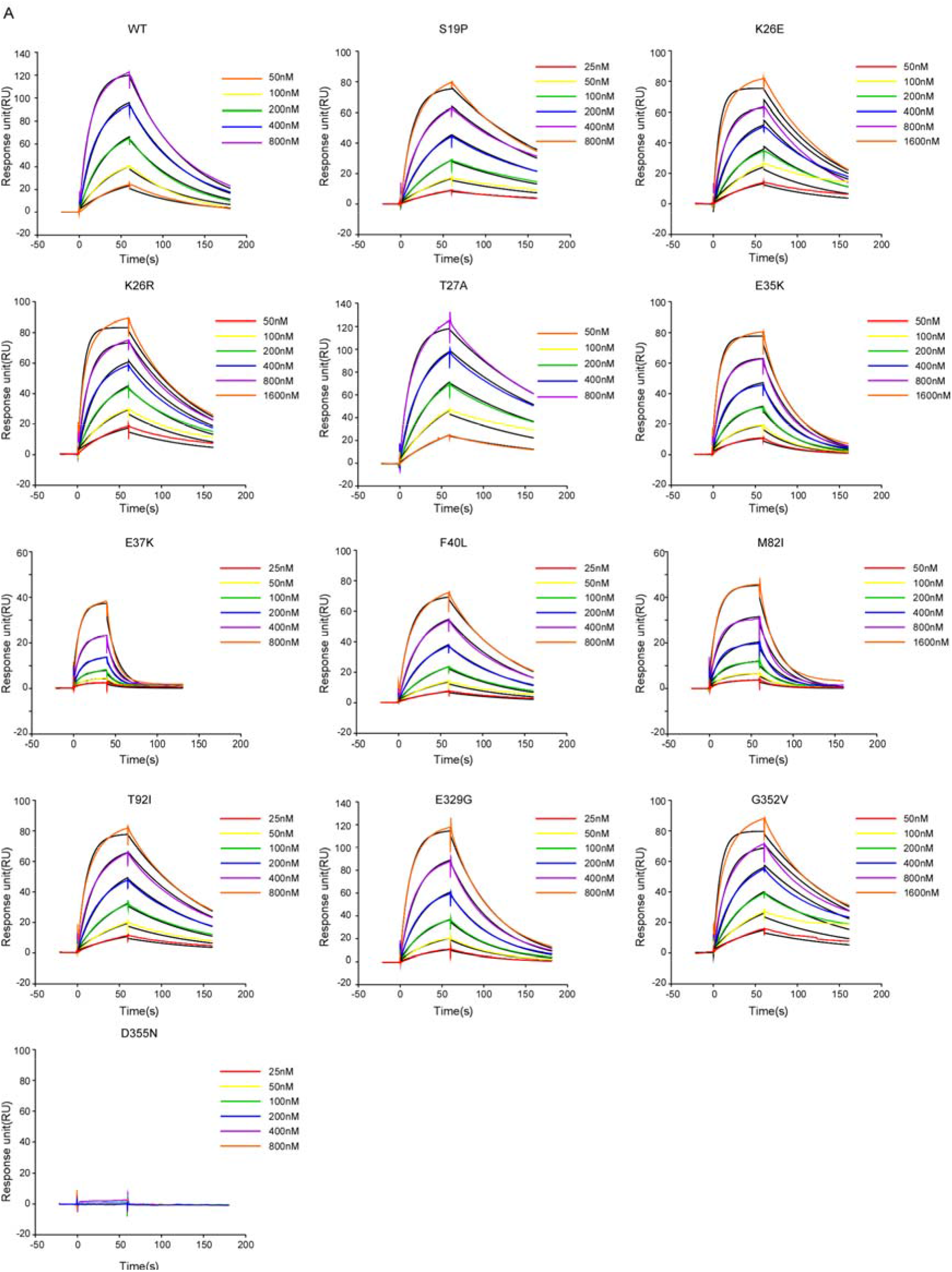
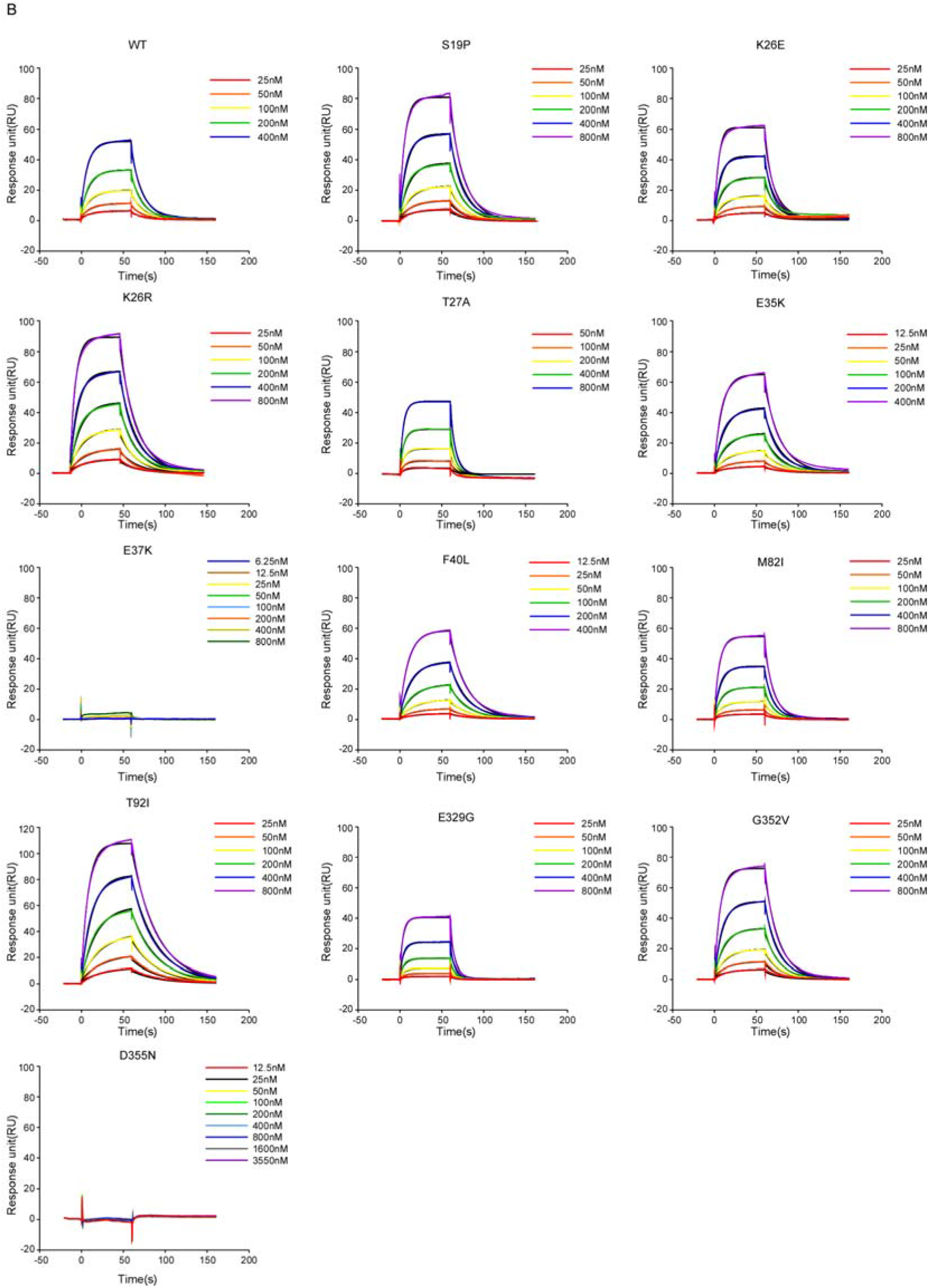
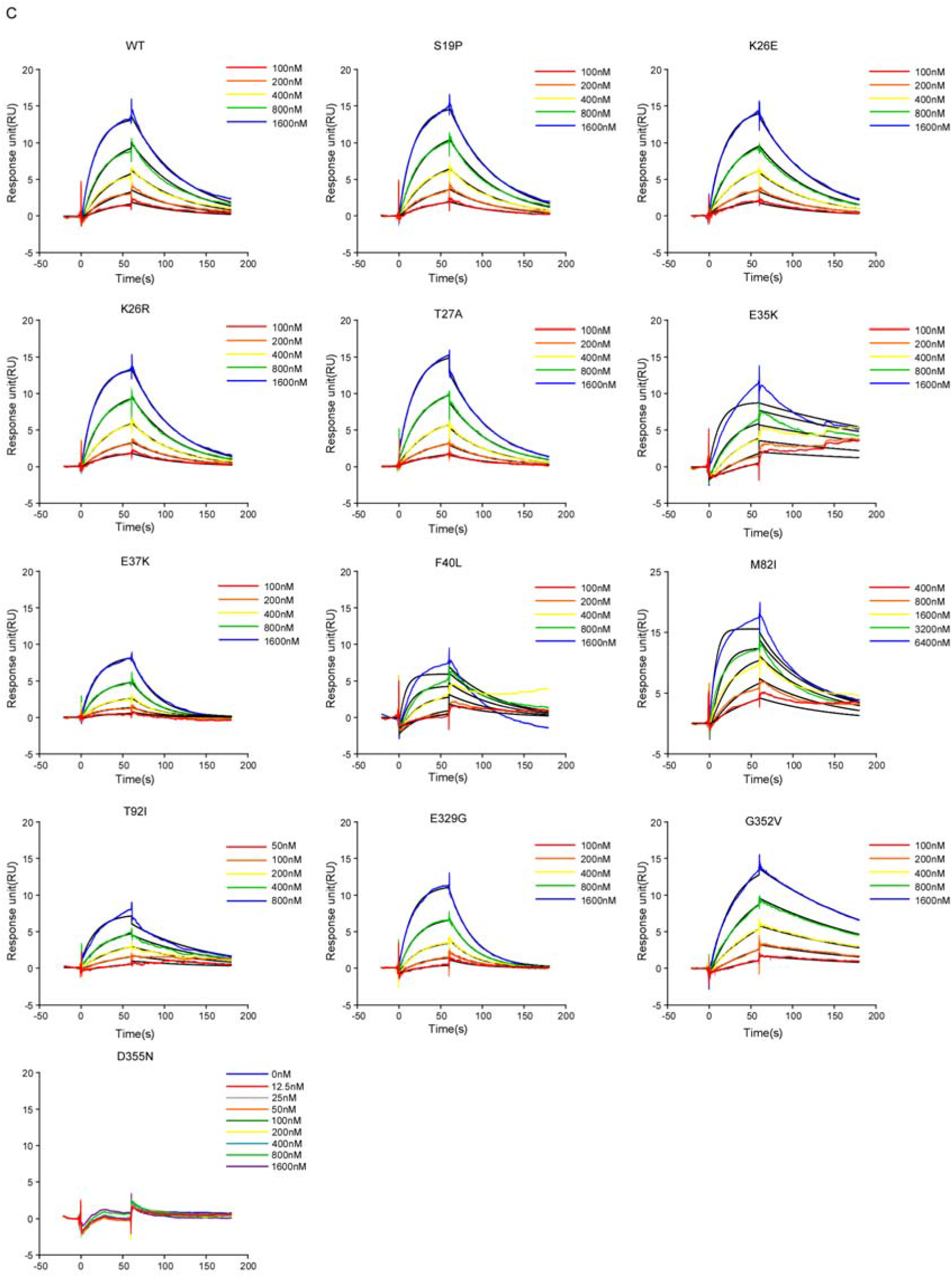
Characterizing the binding affinity of ACE2 variants with coronavirus spike proteins by SPR analysis. SARS-CoV-2 RBD (**A**), SARS-CoV RBD (**B**) or HCoV-NL63 RBD (**C**) was immobilized to a CM5 sensorchip (GE Healthcare); ACE2 SNV proteins of different concentrations were injected. Response units were plotted against protein concentrations. The K_d_ values were calculated by BIAcore 3000 analysis. This experiment was independently repeated three times with similar results and representative results are shown.

